# Experience and behavior modulate piriform cortex odor representation in freely moving mice

**DOI:** 10.1101/2024.02.08.579489

**Authors:** Ian F. Chapman, Martin A. Raymond, Max L. Fletcher

## Abstract

In rodents, activity in the piriform cortex (PC) has been shown to reliably encode the identity of olfactory information within single sessions of odor delivery. However, recent evidence from chronic PC recordings found significant unreliability in this ensemble code over longer periods. The causes of this phenomenon, termed representational drift, are still being investigated across multiple sensory systems, but prior work has suggested a role for animal behavior in this observed unreliability of coding. To explore this possibility in PC, we recorded from the same populations of neurons in freely-moving, awake mice using micro-endoscopic calcium imaging as they gained passive experience with a panel of odorants over 5 consecutive days. As in prior studies, PC odor responses within a single session could be used to accurately decode odor identity. However, responses became less consistent across days of experience as odor-evoked response properties of the neurons shifted with experience. During these recordings, within and across sessions, decreases in olfactory investigative behavior correlated with decreased odor-evoked response from PC neurons. Similarly, decreases in odor investigation correlated with a decrease in representational consistency, and trials with greater odor investigation could be used to decode odor identity from PC neurons more accurately over time. Overall, this data supports recent evidence of long-term shifts in the ensembles of PC neurons encoding odor-identity (drift) but supports a role for behavioral modulation of overall PC activity and ensemble response consistency.

## Introduction

The advent of reliable techniques for longitudinal, single-cell resolution recordings has, in some contexts, demonstrated a surprising unreliability in the long-term encoding of information (Deitch et al., 2021; Driscoll et al., 2017; Marks and Goard, 2021; Schoonover et al., 2021; Ziv et al., 2013). While not readily apparent in all aspects of sensory processing, detailing the presence, impact, and broader circuit implication of this phenomenon, termed representational drift, has become a necessary point of study in many sensory modalities (Driscoll et al., 2022; Rule et al., 2019). In the olfactory system, the first synaptic target after sensory transduction, the olfactory bulb (OB) transforms sensory neuron input into highly reliable and spatially organized patterns of activity that represent the chemical properties of an odor (Bozza et al., 2004; Fletcher et al., 2009; Mori et al., 2006; Uchida et al., 2000; Wachowiak and Cohen, 2001). Longitudinal recordings in the OB have demonstrated passive and associative learning-dependent plasticity but generally show stable representations of chemically distinct odors at both the level of the glomeruli and OB output mitral and tufted (M/T) neurons (Burton et al., 2022; Chu et al., 2016; Gschwend et al., 2015; Kass et al., 2013; Kato et al., 2012; Ross and Fletcher, 2018).

Less is known about the reliability of the odor representation in downstream targets of the OB due to the technical challenges in collecting longitudinal recordings in these regions. Multiple types of neural recordings in humans and animal models have indicated the piriform cortex (PC) as a primary contributor to the formation of odor representations (Gottfried, 2010; Gottfried et al., 2002; Kadohisa and Wilson, 2006; Rennaker et al., 2007; Stettler and Axel, 2009; Wilson, 2000; Wilson and Sullivan, 2011), with broadly distributed ensembles of neurons, particularly in the anterior PC (aPC) able to represent olfactory stimuli identity (Bolding and Franks, 2018, 2017; Kadohisa and Wilson, 2006; Miura et al., 2012; Pashkovski et al., 2020; Rennaker et al., 2007; Roland et al., 2017; Stettler and Axel, 2009), but the stability of this representation within the same population of neurons has only recently been explored across multiple recording sessions. Two recent studies, one leveraging chronic 2-photon calcium imaging (P. Y. Wang et al., 2020) and the other chronic extracellular single-unit recordings (Schoonover et al., 2021), provide the first evidence regarding the consistency of aPC odor representation over time and experience. While both indicate a degree of consistency in aPC odor representations arising from repeated, frequent odor experience, the only study to record over multiple weeks of experience found a surprising lack of consistency across longer time scales (Schoonover et al., 2021).

Work in other sensory systems and animal models has found evidence that behavior may play a role in the observed inconsistency of a stable recording population (Cowley et al., 2020; Liberti et al., 2022; Sadeh and Clopath, 2022). Whether this is the case in aPC remains unclear; prior work has demonstrated both non-associative and associative dependent learning effects in aPC (Barnes et al., 2011; Chapuis and Wilson, 2012; Chen et al., 2011; Meissner-Bernard et al., 2019; Wilson, 2000), but only a few learning paradigms have been implemented with chronic recordings, with minimal impacts found on stimulus coding (Schoonover et al., 2021; P. Y. Wang et al., 2020). Additionally, previous awake recording paradigms restrict animals by head-fixation (Bolding et al., 2020; Bolding and Franks, 2018, 2017; Schoonover et al., 2021; P. Y. Wang et al., 2020) or only present odors at a controlled port (Miura et al., 2012; Poo et al., 2022). This allows for optimal temporal control of odor stimuli but does not represent a natural implementation of the olfactory system and demands behavioral attention from the animal.

We sought to further explore the dynamics of odor coding in aPC over time by recording chronically from the same cell populations and under freely moving conditions using micro-endoscopic 1-photon calcium imaging. To mimic natural odor conditions more closely, mice were placed in a small chamber into which brief, uncued pulses of odorants were presented while video from a side-angle behavior camera was used to generate kinematic measures of animal behavior during the recording. Similar to prior results in head-fixed animals, we find that passive odor presentations can elicit aPC ensemble responses that reliably encode odor identity despite behaviorally correlated changes to overall activity in aPC during each session. Across days, however, we find decreases in response strength similar to effects found in the OB (Kato et al., 2012; Ross and Fletcher, 2018), as well as decreases in ensemble consistency with experience. Importantly, the degree of cross-day consistency in odor response correlated well with the investigatory behavior of the animal, indicating a combined role for behavioral state alongside experience in influencing the reliability of aPC odor representations over time.

## Results

Freely-moving mice were presented with trials of 6 chemically distinct odorants over 5 consecutive days while GCaMP6s fluorescence was recorded from individual neurons (**Figure 1A, C**). Our imaging approach allowed us to collect long-term recordings from a large population of individual neurons across layers 2 and 3 of aPC (**Figure 1B**). To measure odor diffusion within the box during presentations, a photoionization detector (PID) was placed along the back wall of the chamber. We found that odors quickly rise to peak concentration and stay steady throughout the presentation (**Figure 1D**). Animals were not cued for the start of each trial, allowing for natural sampling of odors without forced direction or attention. The odor panel (**Figure 1A**) was not chosen to provide a difficult discriminatory challenge but rather to allow for multiple different odors to potentially evoke activity in the region of aPC in which the gradient index (GRIN) lens was chronically implanted for each animal. Odors were presented in pseudorandomized blocks of trials (10 total for each odor), without any repeated presentations of the same odor given previously noted effects of habituation on OB and aPC odor responses (Chaudhury et al., 2010; Kadohisa and Wilson, 2006; Linster et al., 2009; Wilson, 2000, 1998). Two-dimensional maps of the spatial footprints of neurons generated by GCaMP recordings allowed for reliable tracking of neurons across all days of odor presentations (**Figure S1**). Combined, this enabled us to generate a pooled dataset of 687 individual neurons from 14 mice that we tracked over all 5 sessions of odor experience.

**Figure 1.**
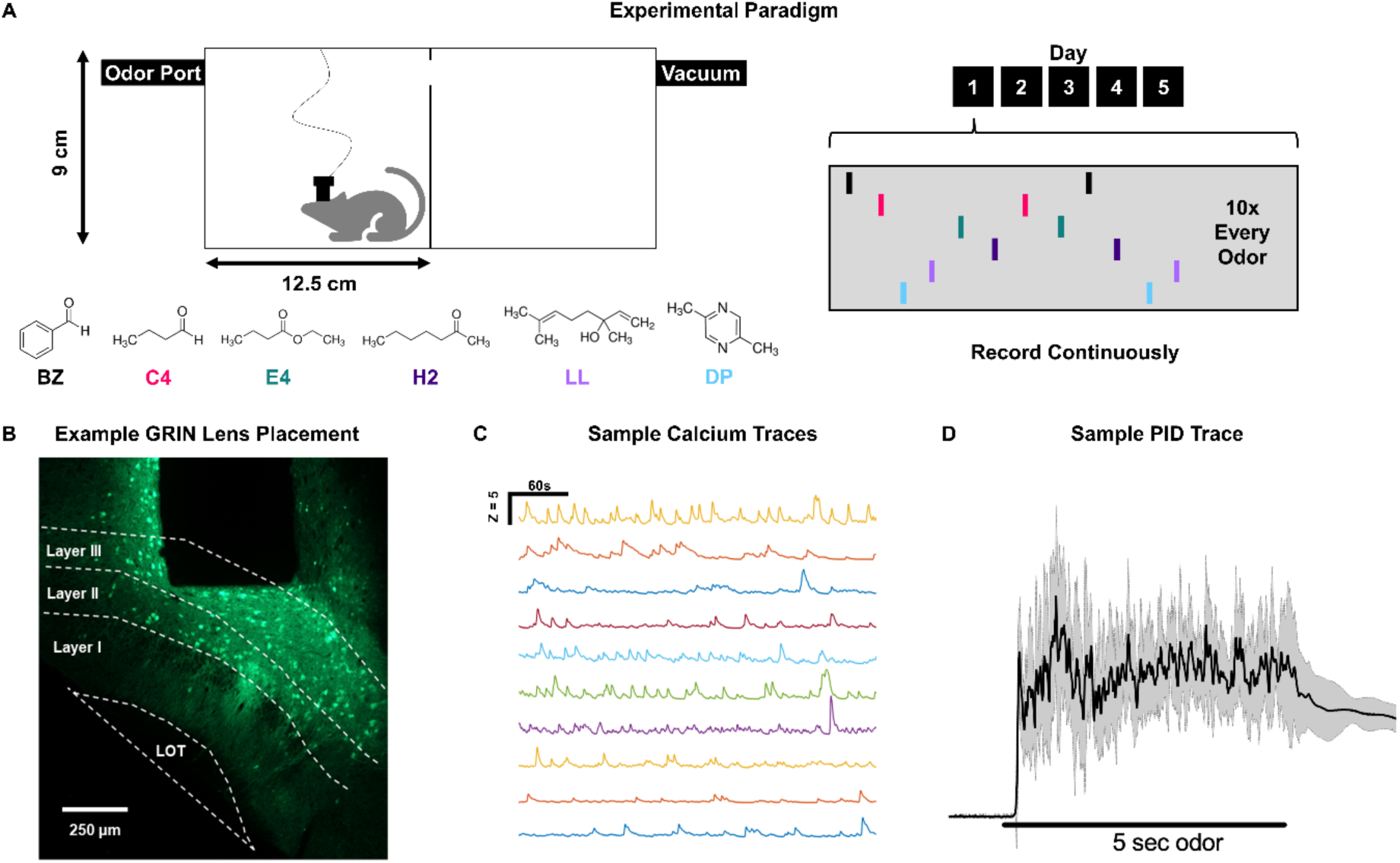
Recording and odor presentation paradigm. **(A)** Schematic of the recording apparatus and paradigm. 10 trials of 6 odorants were presented over 5 consecutive days of passive odor experience while neuronal activity was continuously recorded via miniscope. **(B)** Example GRIN lens implantation site from one animal in aPC; annotated to show the primary layers. **(C)** Example calcium traces from 10 neurons recorded over 400 s at the start of a recording session. **(D)** Mean PID trace (black line, gray bars denote SD) taken from 10 presentations of E4. The PID intake port was placed at the back of the chamber opposite the odor port.

### Isolating odor responses from freely moving animals

At first pass, we measured the maximal response evoked during the odor presentation relative to activity immediately before the odor presentation (ΔF) (Gong et al., 2020; Staszko et al., 2022). Given the fast rise of odor concentration within the box, we expected responses within a single session to be clearly discriminable as in other studies (Bolding et al., 2020; Bolding and Franks, 2018, 2017; Pashkovski et al., 2020; Roland et al., 2017; Schoonover et al., 2021). However, when using a response window spanning the 5 seconds of the odor presentation (similar to previous studies) (Pashkovski et al. 2020; Roland et al. 2017; Wang et al. 2020b), we found that within odor trial-to-trial correlations were exceptionally low (**Figure 2A**) (within odors: μ=0.135 ± 0.006; across odors: μ=0.074 ± 0.002, Mann Whitney test: U = 43311, p<0.001). Additionally, linear classifiers trained on the response vectors (**Figure 2B**) (see methods for details) were unable to reliably decode odor identity to an accuracy expected based on past studies (Bolding et al., 2020; Bolding and Franks, 2018, 2017; Pashkovski et al., 2020; Roland et al., 2017). In exploring the source of this inconsistency, we noted an exceptional amount of activity in the aPC outside of the odor presentation windows (**Figure 2C**). Furthermore, the mean response magnitude of all neurons during odor trials was similar to background activity before the trials (**Figure 2D**), indicating that non-odor-evoked activity during the response windows could be strong enough to interfere with the detection of odor-evoked activity. Additionally, given that the animals in this study were freely moving and no overt, non-olfactory cue was provided signaling odor onset, we expected that behavioral attention to the odor might be delayed despite fast diffusion throughout the box.

**Figure 2.**
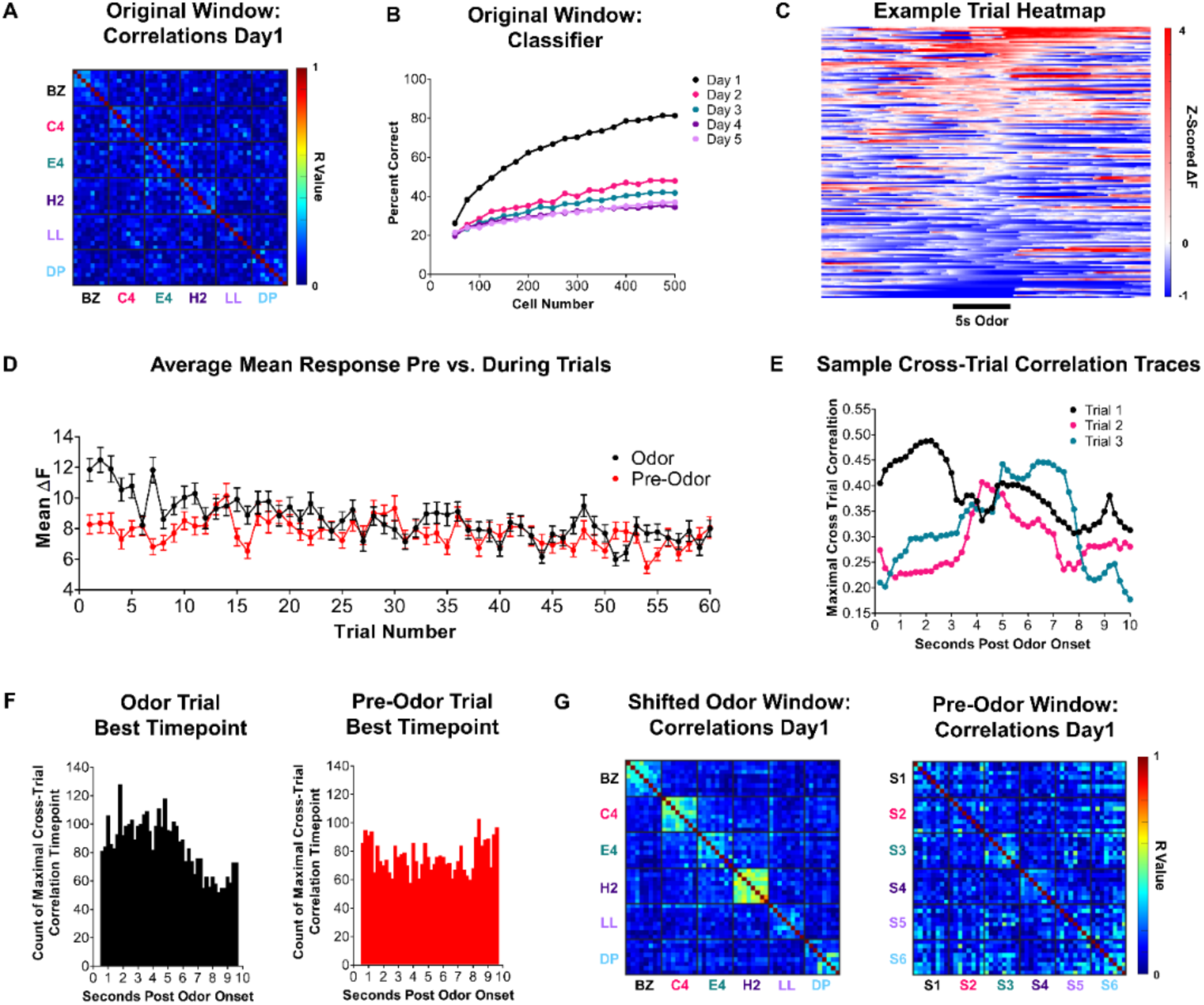
Behavior during odor presentations. **(A)** Trial-to-trial correlations (Pearson’s R) when calculating response values using a 2s pre/5s post window surrounding odor presentation onset. **(B)** Linear classifier (LDA) accuracy in decoding odor identity from response values calculated as in panel A. **(C)** Example heatmap showing activity immediately pre and post-odor presentation for all neurons from one animal (n=154 neurons); neurons were sorted by their maximal response strength during the odor presentation window. **(D)** Average mean ΔF for all neurons at either odor presentation timepoints or in an equally sized window immediately before odor presentation (Pre-Odor); error bars represent SEM, n=792 neurons. **(E)** Representative maximal correlation values of one trial of one odor to the other trials of the same odor when calculating all possible response values in a 10s window post odor onset. (**F)** Distributions of time points with the strongest maximal within odor correlation in a recording session for all time points encompassing the 10s after odor onset (black) or 10s starting a total of 15s before odor onset (red). The first two and last two time points for each histogram were excluded for visualization, given the edge of the effects of the selection window. **(G)** Trial-to-trial correlations after shifting the response timepoint to the maximally correlated timepoint within each odor in the first session (left) or after applying the same method to a window occurring before odor onset of shuffled trials on the same day (right)

Given the number of studies indicating a clear role of aPC in odor perception and representation (Bolding et al., 2020; Bolding and Franks, 2018, 2017; Gottfried, 2010; Gottfried et al., 2002; Kadohisa and Wilson, 2006; Pashkovski et al., 2020; Rennaker et al., 2007; Roland et al., 2017; Stettler and Axel, 2009; Wilson, 2000; Wilson and Sullivan, 2011), we hypothesized that discriminable odor responses were likely embedded in our recordings, but not temporally locked to the response window we initially used due to animal position relative to the odor port, attention to the odor, etc. To test this, we correlated activity from trials of the same odor from a range of all possible time points in a 10-second window after trial onset (see methods for details). We found that, for some trials, peak trial-to-trial response correlation occurred early in the response window (Trial1 in **Figure 2E**), but for other trials, peak similarity in odor response occurred much later in the trial (Trials 2 and 3 in **Figure 2E**). Graphing the distribution of the best-correlated timepoint, we found that, while there was a bias for peak response during the trial, the exact timepoint varied widely after odor presentation onset (**Figure 2F**). This distribution was not found in windows occurring immediately before odor presentation when the same timepoint shifting method was applied (Pre-Odor, **Figure 2F**), indicating that this effect was not an artifact of shifting to a maximally correlated timepoint in aPC activity but was genuinely a result of odor-evoked activity. Given this broad range of the maximally correlated response time, particularly including the first two seconds after the odor trial ended, we chose to shift our window for response measurement to the peak within-odor correlation timepoint found within a 7-second window after trial onset. This shifted response window allowed us to isolate the time points within a recording session where aPC responses best-represented odor identity (within odor correlations: μ=0.325 ± 0.011; across odor correlations: μ=0.114 ± 0.002, Mann Whitney test: U = 11845, p=0.001). (**Figure 2G**). This method does capture a moderate amount of correlated activity from shuffled trials (**Figure 2G**) such that within odor correlations are significantly higher than across odor correlations, albeit with a very small effect size. Since we are aligning trials to their most correlated point of activity, we are necessarily forcing the data to be as similar as possible within the window; this alone could lead to some artificial increase in trial-to-trial correlation. Additionally, however, it is presently unclear if the activity we observed outside of the odor presentations is truly “spontaneous” or reflects PC responses to the animal’s behavior (e.g., grooming, olfactory exploration, etc.); correlated activity in the pre-odor window may be a consequence of coordinated PC activity that reflects some other behavior. Despite these potential drawbacks, this approach more effectively captures odor-evoked information within PC responses under freely-moving conditions (**Figure 2G**). Having isolated odor responses, we address two primary questions through this dataset: how consistent is odor-evoked aPC activity across days of experience? And does animal behavior impact this activity?

### Population response properties

Given past studies indicating broad changes in the overall activity of the olfactory system with odor experience (Barnes et al., 2011; Chapuis and Wilson, 2012; Chen et al., 2011; Kato et al., 2012; Meissner-Bernard et al., 2019; Ross and Fletcher, 2018; Schoonover et al., 2021; Wilson, 2000, 1998), we first explored odor-evoked population activity across our recording sessions. We first compared the mean odor-evoked ΔF of all neurons by averaging across odors for each trial number to assess changes in activity both during and across sessions. Overall, we found a significant decrease in total response magnitude over days (Repeated measures ANOVA: F(3.607, 400.4) = 8.734, p=0.001, post hoc test: Day 1 versus Days 3, 4, and 5, p=0.001) (**Figure 3A**). This effect matches reports of decreases in OB responses over days (Kato et al., 2012; Ross and Fletcher, 2018). Within each day, response magnitude decreased within a session (2-way ANOVA: main effect of trial, F (4.971, 323.1) = 22.78, p=0.001, Trial 1 versus Trial 8 post hoc tests, p≤0.024 for each day). This within-session decrease rebounded to initial levels on each subsequent day as no significant differences were seen in response magnitudes of the first trial of each odor between days (2-way ANOVA: main effect of day, F (4, 65) = 0.651) (**Figure 3B**).

**Figure 3.**
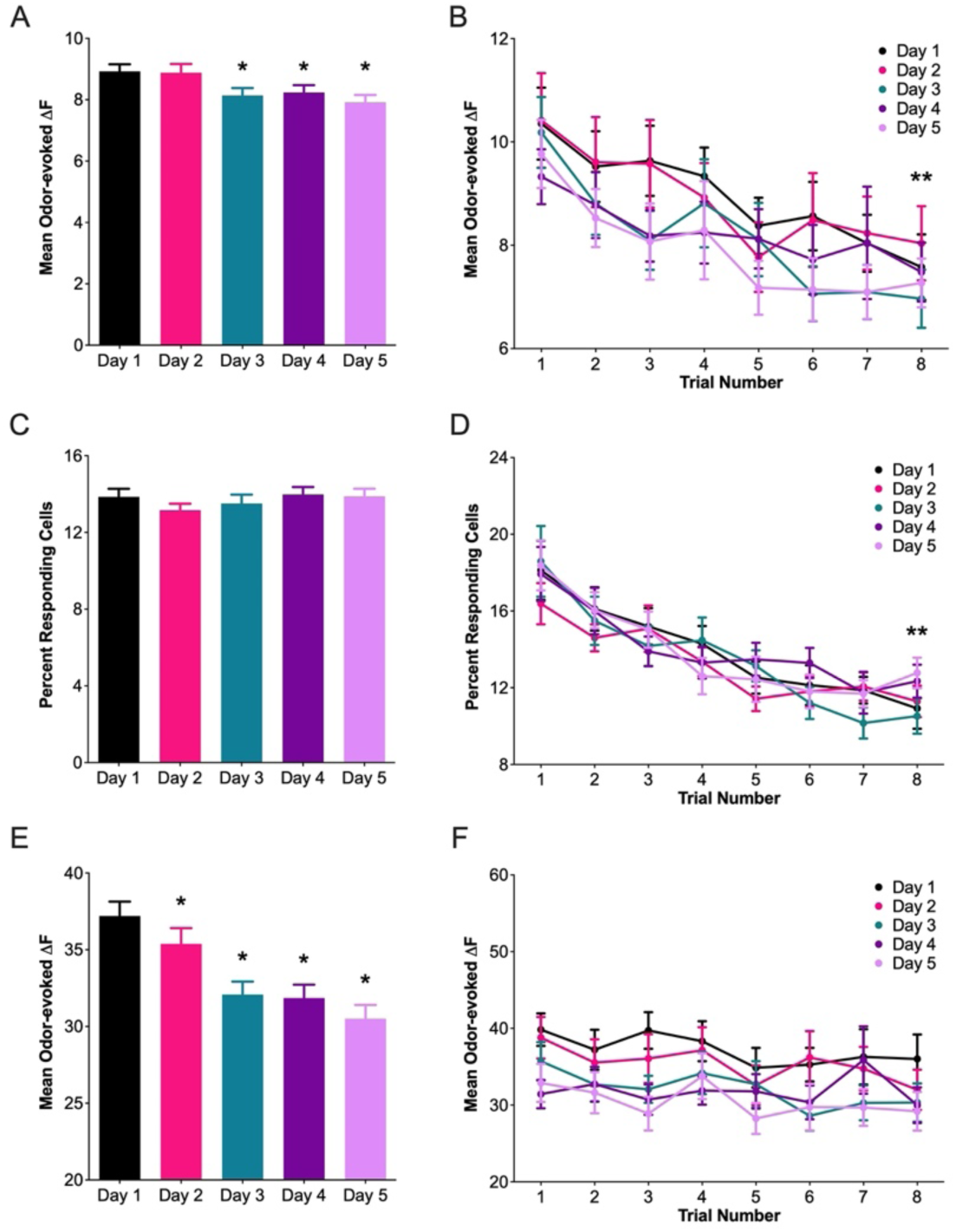
Population response properties during and across sessions. Odor-averaged measures of activity in aPC neurons during odor presentations; or all graphs, each datapoint represents mean, and error bars represent SEM calculated across animals; n=687 neurons from n=14 animals. **(A)** Odor-evoked ΔF from all neurons averaged from all trials on a given day or (**B)** separated by trial before averaging across odors. **(C)** Percent of recorded neurons responding to a trial averaged from all trials on a given day or **(D)** separated by trial number before averaging across odors. (**E)** Odor-evoked ΔF from only significantly responsive neurons (> 2.5 s.d. increased response and z-score > 1) averaged from all trials on a given day or **(F)** separated by trial before averaging across odors. * denotes a significant difference from Day 1. ** denotes a significant difference from Trial 1.

Prior studies have indicated that only a percentage of aPC is activated by any given odor (Bolding et al., 2020; Bolding and Franks, 2017; Pashkovski et al., 2020; Roland et al., 2017; Schoonover et al., 2021; Stettler and Axel, 2009), so we next investigated population activity of neurons that were significantly responsive on a given trial. Though difficult to compare response magnitude across recording techniques and states of anesthesia, the percent of neurons responding to a given odor in our data is similar to other studies (Bolding et al., 2020; Bolding and Franks, 2018, 2017; Pashkovski et al., 2020; Roland et al., 2017; Schoonover et al., 2021; Stettler and Axel, 2009). While the percent of neurons responding to odor was stable across days (Repeated measures ANOVA: F(3.801, 421.9) = 1.439, p=0.222) (**Figure 3C**), the percent of responsive neurons did drop across trials within each day (2-way ANOVA: main effect of trial, F (1.939, 25.20) = 19.55, p=0.001, Trial 1 versus Trial 8 post hoc tests, p≤0.004 for each day) (**Figure 3D**). The mean magnitude of the significantly responsive neurons also decreased across days (Repeated measures ANOVA: F(3.758, 417.1) = 21.65, p=0.001, post hoc test: Day 1 versus Days 2-5, p=0.001) (**Figure 3E**). However, this population did not show significant decreases in magnitude within sessions (2-way ANOVA: main effect of trial, F(4, 64) = 1.53, p=0.204) (**Figure 3F**). This indicates that, despite experience-dependent changes in population response dynamics, we still found populations of neurons that responded robustly throughout recording sessions to odor presentation.

### Odor coding within days

Given that we found changes in overall aPC activity within days, we next asked what effect this had on the encoding of odor information. We first calculated pairwise ensemble correlations using the odor-evoked response of all neurons to each odor within each session (**Figure 4A**). This recapitulates analyses performed in acute recording studies, treating each session as an independent point of analysis (Bolding et al., 2020; Bolding and Franks, 2018, 2017; Pashkovski et al., 2020; Roland et al., 2017). Correlations of trials of the same odor were clearly separable from correlations of trials across odors, an expected result given the distinct structural differences between odors selected in our panel. Overall, when correlations were averaged across odors, we found a significant decrease in within-odor correlation across days (Repeated measures ANOVA: F(1.491, 249.0) = 375.4, p=0.001, post hoc test: Day 1 versus Day 5, p=0.001) (**Figure 4C**) suggesting that trial-to-trial response stability degrades over time.

**Figure 4.**
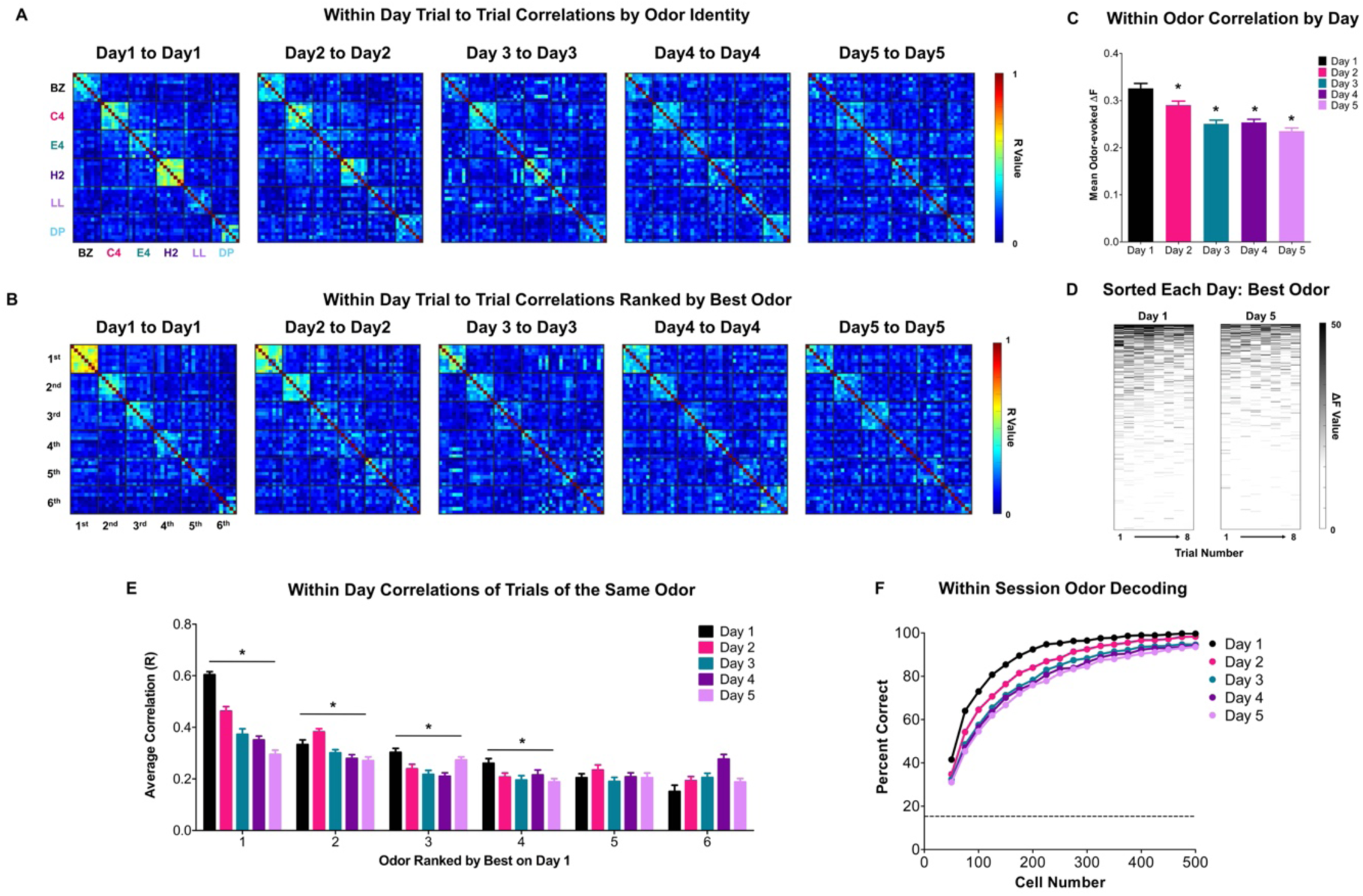
Reliable odor coding within sessions of freely-moving odor experience. **(A)** Trial-to-trial correlation (Pearson’s R) matrices generated between all trials for all neurons within a day of recording, organized by odor identity. (**B)** Same as in panel A, but after reorganizing for the most strongly correlated odor on Day 1 (“best”) for each animal instead of by odor identity before generating the correlation matrix. **(C)** Average R-value for all within-odor comparisons from panel B (trials of BZ to BZ, trials of C4 to C4, etc.) by day; mean and SEM shown. **(D)** Heatmaps showing the responses from all 687 neurons either on Day 1 or Day 5 to the best correlated odor; sorting each day by mean response strength. **(E)** Quantification of the within odor correlation values from panel B; correlations shown from within each Day, with odors organized by the “best” odor on Day 1. Each bar represents the mean of all unique trial-to-trial correlations, and error bars denote SEM. * denotes a significant difference between Days 1 and 5. **(F)** Odor decoding from increasingly large random subsets of the neural data using an LDA classifier; each classifier trained and tested on data from the same day using leave-one-out validation. For each data point, 100 iterations of randomly sampled neurons from the dataset of size n=neuron number were used to train each model. The same neurons were used across days at each iteration for consistency.

However, within the session, mean odor correlations were not equivalent for each odor, with most animals showing a preferential response to a subset of odors from the panel, particularly 2-heptanone (**Figure S2**). While there is abundant evidence of broad distribution of olfactory bulb inputs to aPC (Haberly and Price, 1978; Miyamichi et al., 2011; Neville and Haberly, 2004; Sosulski et al., 2011) and previous studies have found broadly distributed ensembles of neurons in PC that encode odor information (Bolding and Franks, 2018, 2017; Pashkovski et al., 2020; Roland et al., 2017; Stettler and Axel, 2009), our odor presentation paradigm diverges significantly from past studies with fixed odor presentation. Increased variability in animal behavior and odor turbulence before reaching the nose may only result in consistent activation of a smaller subset of strongly connected neurons within the OB to PC circuit. We, therefore, hypothesized that this differential responsivity may affect changes to within-day ensemble correlations; odors that only weakly activate aPC in a given animal may show much weaker consistency over time, whereas more information may have been retained from strongly activated odors. To test this, we first ranked each animal’s odor responses by the strength of their Day 1 mean trial-to-trial correlations, and all odors were re-sorted each day based on that “best” odor ranking (**Figure 4B**). Using a 2-way ANOVA of data from all days, we found a main effect of odor rank (F(1.954, 52.76) = 1508, p=0.001), indicating that responses to the “best” ranked odors were significantly more consistent. When pooled across days, post hoc tests revealed that mean day-to-day correlations of the best odor were significantly stronger than those of all other odors (p=0.001). When comparing day-to-day changes for each odor rank, we found a main effect of the day (F(2.252, 60.79) = 488.6, p=0.001), suggesting that, on average, trial-to-trial correlations decrease over days (**Figure 4E**). At the level of individual odors, post hoc tests revealed that the effect was significant for the top four odors (Day 1 versus Day 5, p=0.001) (**Figure 4E**), indicating that response consistency decreased even in the most strongly represented odor for each animal (**Figure 4D, E**).

To determine whether this decrease in response stability impacted odor coding, we trained a linear classifier on responses from increasingly larger random samples of neurons from each day to test how accurately odor identity could be decoded within each session. On later days, the linear classifier required more neurons from the dataset to correctly decode odor identity from the neural data (**Figure 4F**), but activity within any day of recording contained ample information to reach or approach 100% accuracy in decoding odor identity. Taken together, this indicates that, while the density of information may decrease with experience, potentially as a result of the previously noted changes in response strength (**Figure 3E**), aPC responses on any given day are adequate for correctly identifying odor identity within that day despite a reduction in trial-to-trial correlations.

### Odor coding across days

We next investigated response stability across days in our paradigm. To explore this, we compared response correlations for all trials on each day to Day 1 and found a significant lack of consistency in ensemble responses to the same odor (**Figure 5A**). When accounting for the previously noted uneven distribution of strongly activated odors, we still found a decrease in response consistency for all odors, including the best odor (**Figure 5B, C**). Similar to the decrease found when previously assessing consistency within sessions (**Figure 4E**), we found an overall decrease in response similarity across days (2-way ANOVA, the main effect of the day: F(4, 1464) = 492.9, p=0.001). Post hoc tests revealed this effect was significant for all odors individually (p=0.001) (**Figure 5D**). Importantly, for all odors and days, this decrease in consistency was notably stronger when looking across sessions (each day compared to the previous day) compared to within sessions (2-way ANOVA, F(1, 2200) = 291.8, p=0.001, within versus across post hoc test for each day: p=0.001). (**Figure 5E**). This decrease in cross-day response consistency led to worse performance of linear classifiers in decoding odor identity on each day when models were only trained to the first day of the dataset (**Figure 5F**). Notably, these classifiers still performed well above chance level, indicating some conservation of odor information, but certainly not a 1:1 maintenance of representation across days within the same population. Additionally, classifier accuracy on Days 2-5 began to asymptote at well below 100% accuracy, indicating the possibility that, even if we could expand the size of our collected dataset, a fundamental change in the odor response profile of at least some of the neurons may have occurred across sessions.

**Figure 5.**
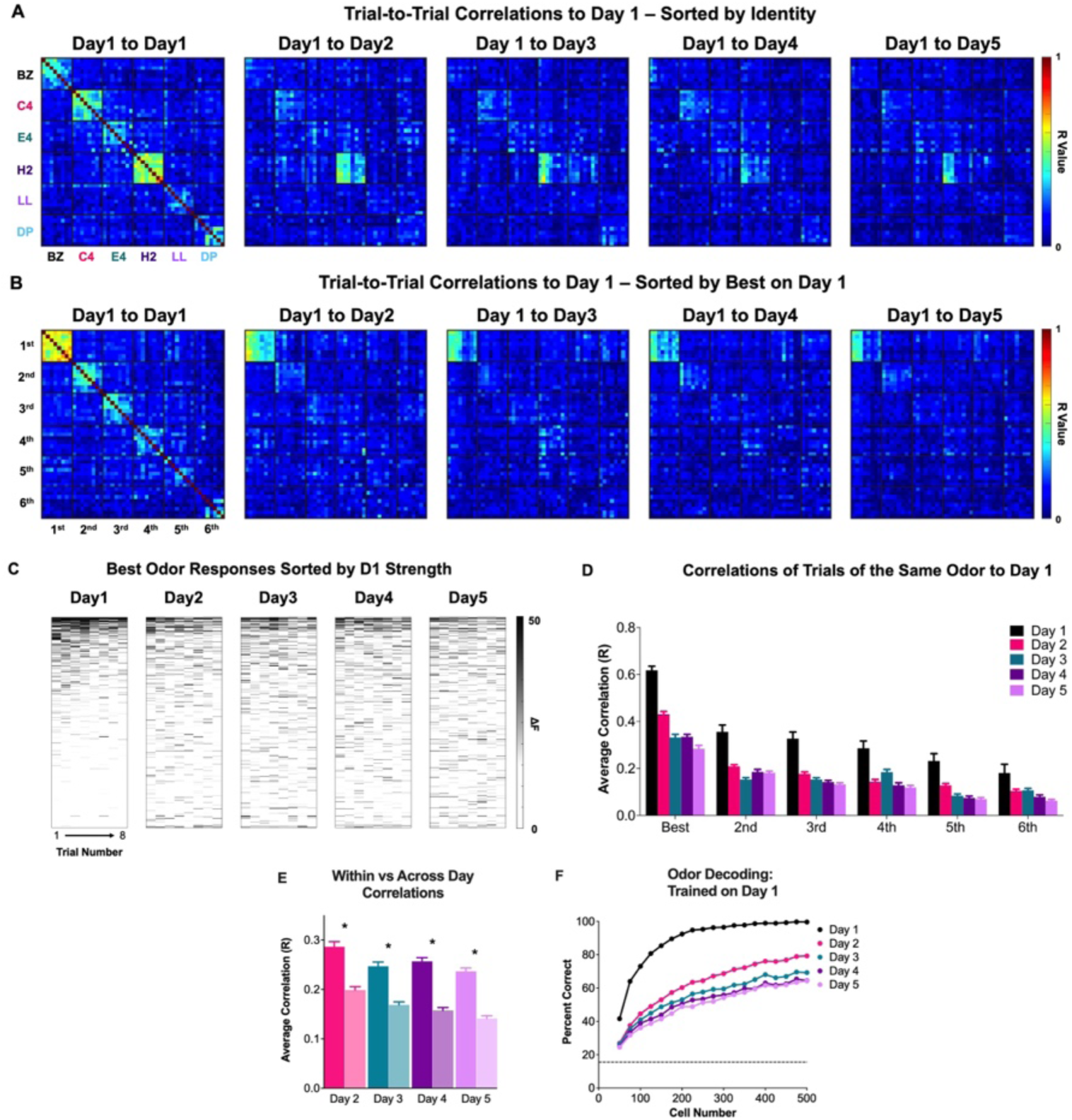
Decreased consistency in cross-session odor coding. **(A)** Trial-to-trial correlation (Pearson’s R) matrices generated by comparing all trials of a given day to all trials on Day 1, organized by odor identity. **(B)** Same as in panel A, but after reorganizing for the most strongly correlated odor on Day 1 (“best”) for each animal instead of by odor identity before generating the correlation matrix. **(C)** Heatmaps showing the responses from all 687 neurons on Days 1-5 to their best odor on Day 1, sorting each neuron for all days by their mean response strength on Day 1. **(D)** Quantification of the within odor correlation values from panel B; correlations to Day 1 with odors organized by their best odor on Day 1. Each bar represents the mean of all unique trial-to-trial correlations, and error bars denote SEM. **(E)** Comparison of within-session correlations (from 4E) to correlations to Day 1 values (panel D) for Days 2-5. Each bar represents the mean of all unique trial-to-trial correlations, and error bars denote SEM. * denotes a significant difference. **(F)** Odor decoding from increasingly large random subsets of the neural data using an LDA classifier; each classifier trained on trials from Day 1 and tested on a trial from the given day. For each data point, 100 iterations of randomly sampled neurons from the dataset of size n=neuron number were used to train each model. The same neurons were used across days at each iteration for consistency.

Given recent evidence of representational drift in PC odor responses (Schoonover et al., 2021), we next investigated whether our cross-day changes in odor coding were a result of a reorganization of odor response profiles. Since only a few odors per animal evoked consistent cross-day responses (**Figure 5B**) and only approximately 10-20% of aPC neurons responded significantly to odor presentations (**Figure 3C, D**), there remained a possibility that a subpopulation of consistently responding neurons for those odors could provide more consistent representations of odor identity across days. This would imply exceptionally sparse encoding of odor identity but remains a possibility as we can only sample a small fraction of the total population of aPC from a given animal. To investigate this, we first looked at the specificity of an individual neuron’s responses. We used lifetime sparseness, a method that assesses how broadly tuned a cell’s response is to a panel of stimuli (Rolls and Tovee, 1995). A value of 1 indicates that a cell exclusively responds to a single stimulus, whereas a value of 0 indicates that a cell does not respond differentially to any of the stimuli. Low lifetime sparseness values, therefore, indicate neurons that either respond to no stimuli or are broadly tuned to all stimuli, while high sparseness values indicate that a cell is narrowly tuned to fewer stimuli.

Our average lifetime sparseness values **(Figure 6A)** are lower than previously reported in anesthetized studies (Poo and Isaacson, 2009) and are more in line with values reported from prior awake recordings (Bolding and Franks, 2017; Miura et al., 2012; Schoonover et al., 2021). A small but significant decrease in mean lifetime sparseness was found across days at the population level (Day1 µ = 0.369, Day 5 µ = 0.319, ANOVA F (4, 2744) = 13.26, p < 0.0001) and could be a result of the decrease in odor response magnitude previously observed (**Figure 3E**). Despite this change in the mean, the shape of the distribution remained similar across days, with a small sup-population of neurons showing more narrowly tuned response profiles (**Figure 6A**). Within each day, responses from narrowly tuned neurons could be used to generate classifiers that reach high accuracy much faster than broadly tuned neurons (**Figure 6 B, C**). This is an expected result, as neurons with narrower response values would, by definition, provide more discriminable information for linear classification methods. Nonetheless, this demonstrates that these neurons drove reliable encoding of odor information within a session. We, therefore, tested whether the more narrowly tuned neurons provided more reliable odor information across sessions, but similarly to our whole population data, we still found decreases in cross-session decoding accuracy (**Figure 6D**).

**Figure 6.**
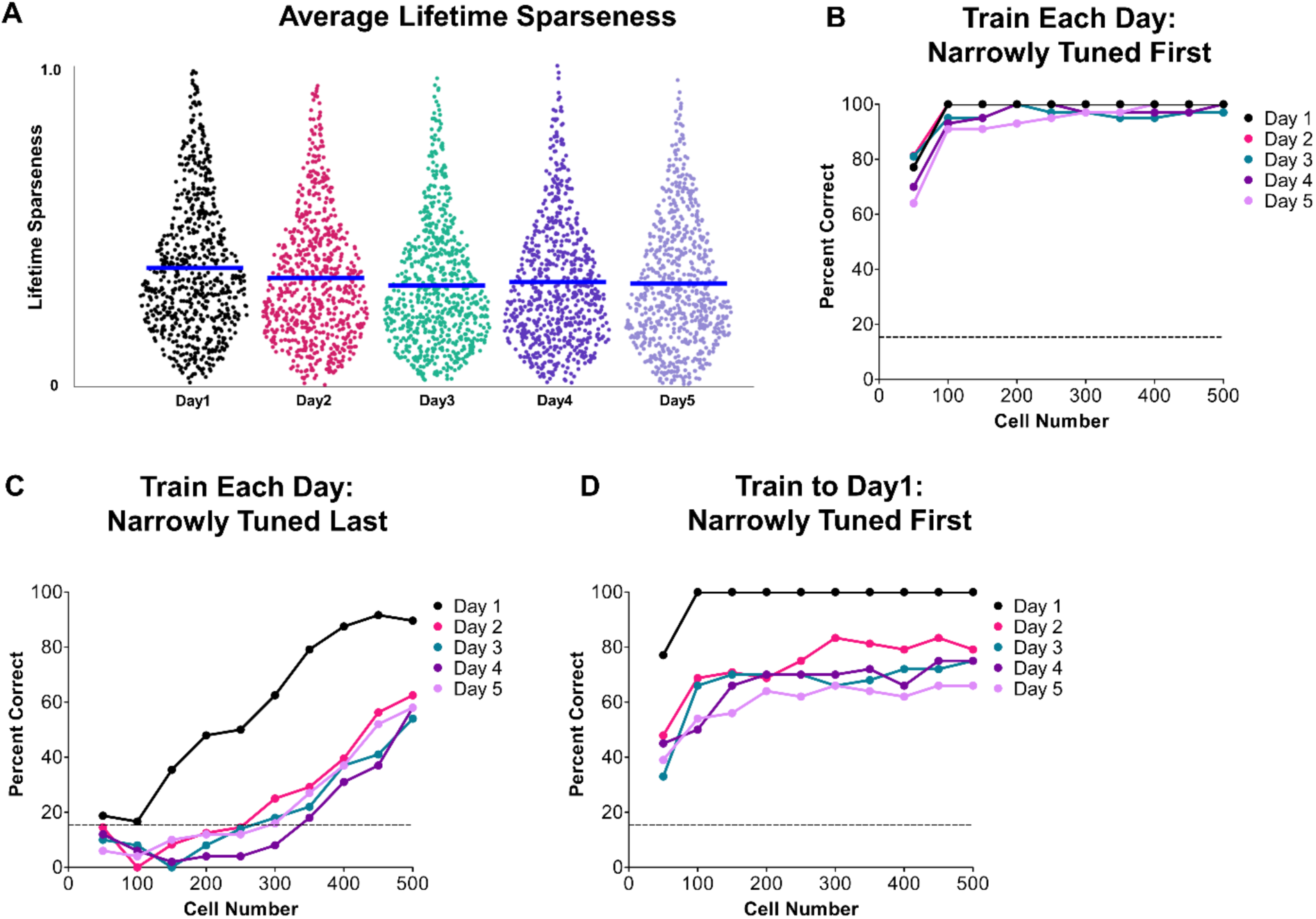
Narrowly tuned sub-populations robustly encode odor identity but are less reliable in encoding identity across days. **(A)** Lifetime sparseness of all neurons calculated each day; mean represented by a blue bar over the distribution of each group. **(B)** LDA classifier results when training/testing with the n=neuron number of most narrowly tuned neurons on a given day using sub-populations of those neurons of increasing size. **(C)** Same classifier method as in panel B, but neurons with the most widely tuned responses were selected first. **(D)** LDA classifier results when only training the models using the most narrowly tuned neurons on Day 1 and testing the model using responses from the same sub-population on subsequent days.

To further characterize why this more robust odor-encoding subset of neurons was still less accurate over time, we explored the response properties of these narrowly tuned neurons to their preferred odor. The preferred odor for each cell was defined as the odor evoking the maximal mean response from a cell on a given day. Responses from all cells with the same odor were pooled and averaged. Using a 2-way ANOVA, we found an interaction of group and odor (F (25, 660) = 18.41, p=0.001). However, no differences in odor response magnitude were found between any best odor group (all odor responses pooled) (main effect of group: F (5, 132)=1.632, p=0.156) or between any of the responses to each of the odors when pooled (main effect of odor: F (4.486, 592.2) = 0.384, p=0.841) suggesting that the tuning curves of narrowly tuned neurons are consistent across odors. As expected, post hoc tests revealed responses of narrowly tuned neurons to their preferred odor are significantly stronger than to the other odors in the panel when sorted on Day1 (BZ best: BZ vs. all p =0.001 or lower, C4 best: C4 vs. all p = 0.001 or lower, E4 best: E4 vs. all p = 0.015 or lower, H2 best: H2 vs. all p = 0.001 or lower, LL best, LL vs all p = 0.001 or lower, DP best” DP vs all p = 0.001 or lower) (**Figure 7A**).

**Figure 7.**
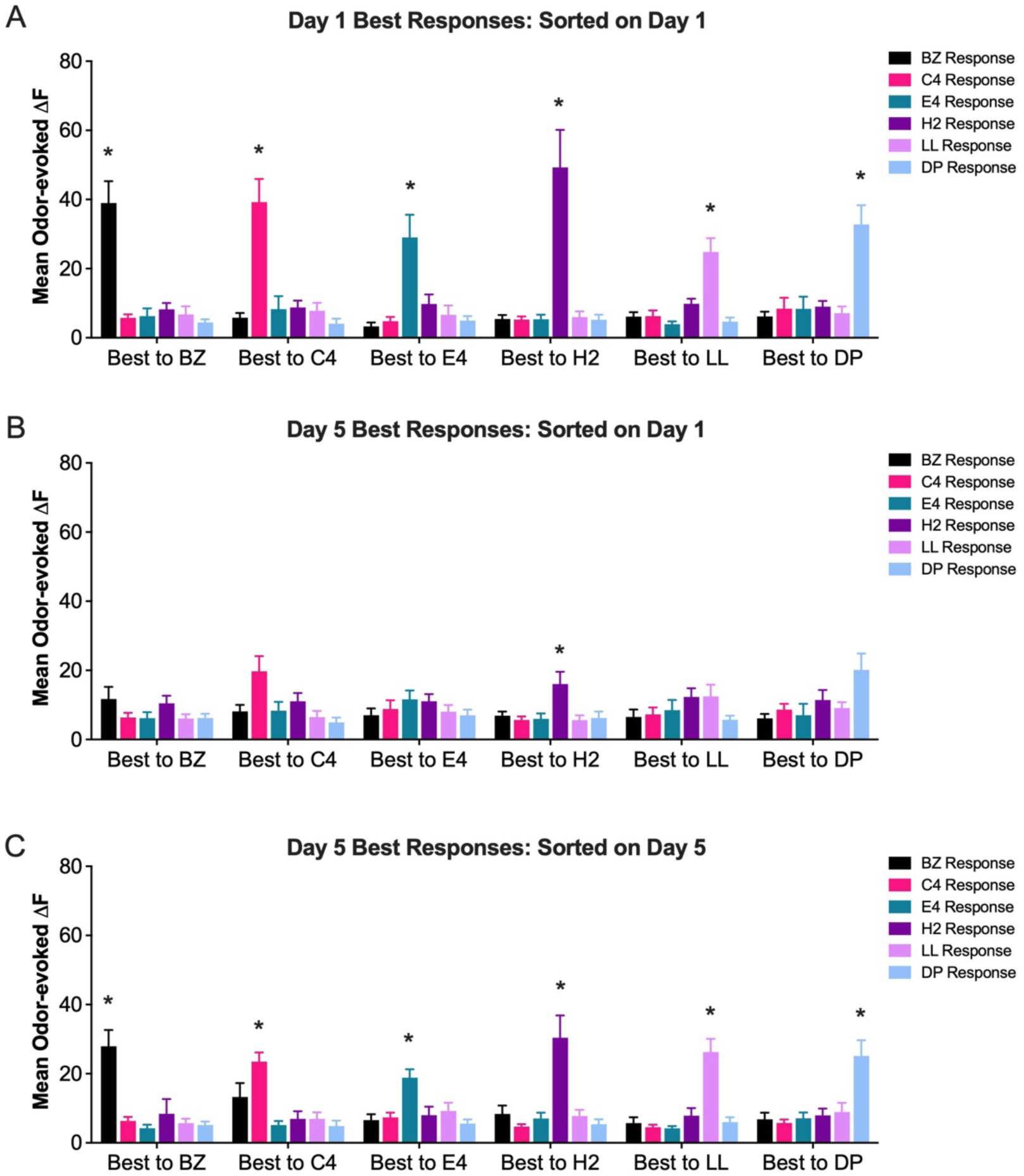
The odor response preference of narrowly tuned neurons is not consistent across days. In all cases, the top 20% of narrowly tuned neurons were selected, and mean-evoked ΔF and SEM were graphed for each sorted best responding sub-population. **(A)** Neurons selected by their tuning on Day 1 and sorted by their odor with the strongest average odor response (best) on Day 1. **(B)** Day 5 responses from neurons selected by their tuning on Day1 (same subset of neurons as in panel A) and sorted by their strongest average odor response (Best) on Day1 **(C)** Neurons selected by their tuning on Day 5 and sorted by their strongest average odor response Day 5. * denote significant difference from all other odor responses.

However, when we sorted day 5 neurons by their preferentially tuned odor on Day 1, we found a decrease in preferred odor response magnitude for all odors except LL (2-way ANOVA, main effect of odor, F (1, 130) = 58.19, p=0.001, post hoc tests: p=0.034 or lower) (**Figure 7B**). As above, we used a 2-way ANOVA to compare the response magnitude of the preferred odor to responses to the other odors. We found an interaction of group and odor (F (25, 660)=2.97, p=0.001), but no differences between any best odor group (all odor responses pooled) (main effect of group: F (5, 132)=1.32, p=0.258) or between any of the responses to each of the odors when pooled (main effect of odor: F (4.515, 595.9)=1.205, p=0.306) (**Figure 7B**). The significant decrease in response magnitude of the preferred odor led to these responses not being significantly different from the responses to the other odors for some of the best odor groups (post hoc tests, p>0.05). However, responses from the preferred odors C4, H2, and DP remained significantly higher in some cases (p<0.045 or lower). Particular in the case of 2-heptanone (H2 vs all other odors, p = 0.001 or lower), this was an expected result as this odor maintained the strongest cross-day response correlation values (**Figure 5A**), and our classifier results indicated at least some conservation of odor information above chance level (**Figure 5F**).

Overall, these findings suggest that the odor preference of individual neurons is not stable across days. However, it is possible that this effect of a loss of odor preference could have simply been the result of the decreased magnitude of strong odor responses previously noted at the population level (**Figure 3E**). This does not appear to be the case as neurons can be resorted by their Day 5 preferences and display similar tuning profiles and response magnitudes compared to Day 1 sorting (**Figure 7C**). While there is some slight overall decrease in preferred odor response magnitude compared to Day 1 (2-way ANOVA, main effect of day, F (1, 264) = 8.47, p=0.004), post hoc tests revealed this decrease is only significant for the H2 preferred group (p=0.025). As above, we used a 2-way ANOVA to compare the response magnitude of the preferred odor to responses to the other odors. We found an interaction of group and odor (F (25, 660)=19.40, p=0.001), but no differences between any best odor group (all odor responses pooled) (main effect of group: F (5, 132) = 1.17, p=0.356) or between any of the responses to each of the odors when pooled (main effect of odor: F (4.486, 592.2)=0.384, p=0.841). As expected, post hoc tests revealed responses of narrowly tuned neurons to their preferred odor are significantly stronger than to the other odors in the panel when sorted on Day1 (BZ best: BZ vs. all p =0.001 or lower, C4 best: C4 vs. all p = 0.014 or lower, E4 best: E4 vs. all p = 0.006 or lower, H2 best: H2 vs all p = 0.002 or lower, LL best, LL vs all p = 0.001 or lower, DP best” DP vs all p = 0.002 or lower) (**Figure 7C**). This dramatic change in response tuning, even within neurons that more robustly represent odor identity, demonstrates that our observed changes in odor coding were not merely the result of decreased signal or increased noise within the aPC responses but rather represented a change in response profile within the population of neurons most sensitive to odor identity. While recorded from a different paradigm and different time scale than prior work (Schoonover et al., 2021), this indicates a similar phenomenon of reorganization of the specific ensembles of neurons representing odor identity in PC over time.

### Impact of behavior on aPC odor coding

Finally, given that our animals were freely moving during our recordings, we wanted to assess whether differential behavior could explain any components of our observed changes in aPC activity and odor response profiles. We quantified investigatory behavior using video recorded from a side-view camera positioned close to the chamber. Using these recordings, we trained a DeepLabCut (Mathis et al., 2018; Nath et al., 2019) model to estimate the position of the ears, nose, body, rear, and body of the miniscope. From this, we generated metrics for odor investigation based on a motion index and stereotypical raising of the nose (Ogg et al., 2018) (**Figure 8A**). To account for any trials where either metric did not meet our confidence criteria (see methods), we combined normalized values for each metric into a single combined behavioral score for each trial that approximated the relative investigation of the odor during the trial.

**Figure 8.**
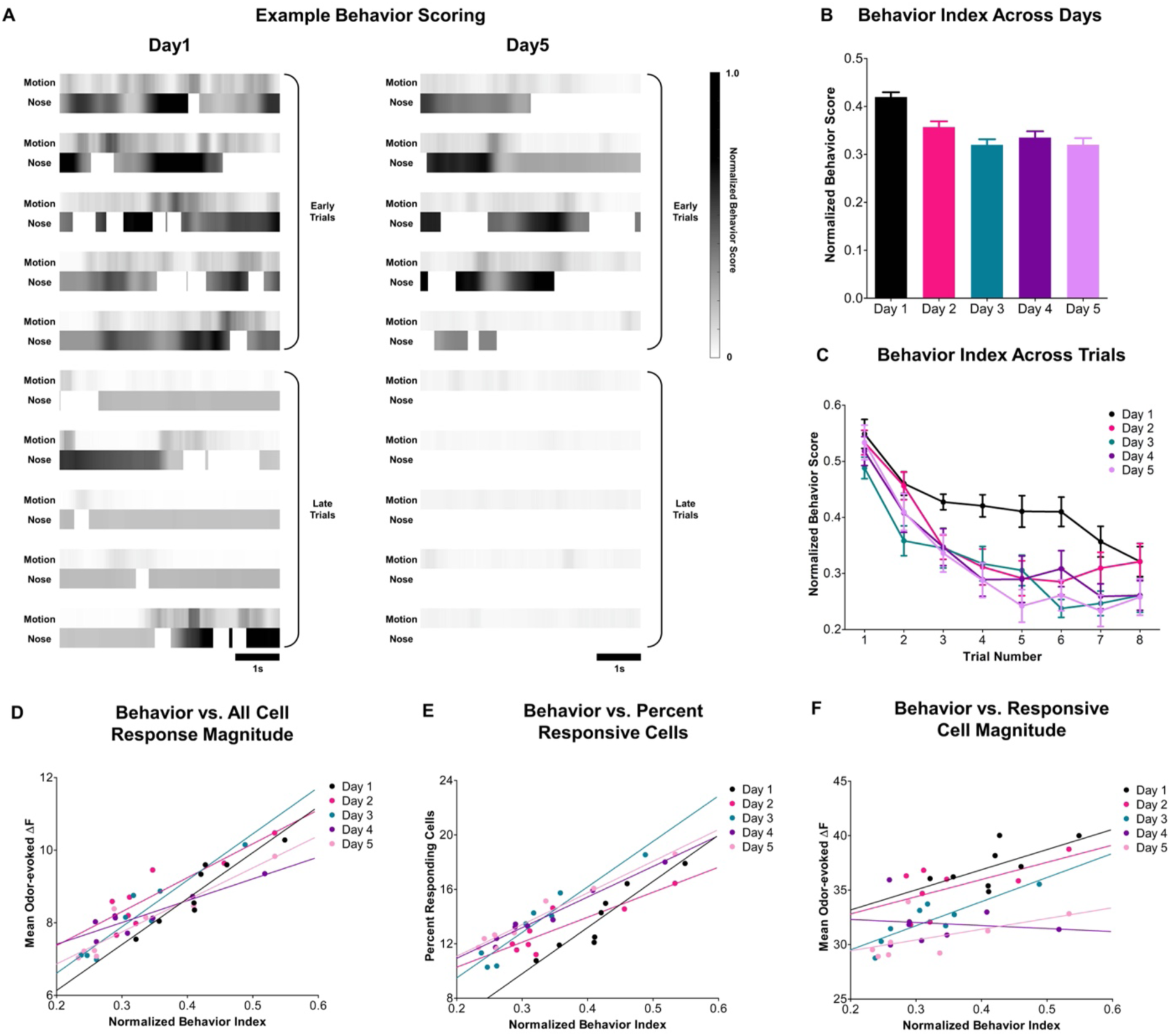
Investigative behavior correlates with aPC response profile. **(A)** Representative heatmaps showing behavior values for either the motion index or raising of the nose during the 5s window used for neuronal response quantification; normalized for the maximum value each of metric for the sample animal for visualization purposes. **(B)** Normalized behavior index values across all trials of a given day for all animals (n=15); mean and SEM shown. **(C)** Same behavior index values as in B, but odor-averaged and separated by trial number; mean and SEM shown. **(D-F)** Odor averaged behavior index scores across trials from C compared to the response magnitude of all cells (Figure 3B), percent of responsive cells (Figure 3D), or response magnitude of significantly active cells (Figure 3F). In all cases, comparisons are made using Pearson’s correlation between the two sets of values within a day; linear regression lines of best fit are shown for each day.

To assess changes in investigation both within and across sessions, we averaged the behavior index across odors at each trial number. Average investigation of the odors is significantly higher on the initial day of odor presentations when the odors are novel (Repeated measures ANOVA, F(3.755, 443.1) = 20.39, p=0.001, post hoc tests: Day 1 versus every other day, p≤0.001 for all comparisons) (**Figure 8B**). Despite this effect, we find that, even during the first session of recording, animals behaviorally habituate to odor presentations, becoming less active on subsequent trials of the odors (2-way ANOVA, F(3.999, 55.98) = 43.14, p=0.001) (**Figure 8C**). The overall larger mean behavior activity score on the first day is driven by a flatter habituation curve within the session, with more investigation occurring on trials 3-7 compared to other days (**Figure 8C**). Interestingly, no significant differences were observed in investigation scores for the first trial of each day (day versus day, Tukey multiple comparisons tests: p<u>≥</u>0.304), suggesting that this habituation recovers between sessions and mice are once again interested in the odors (**Figure 8C**).

To explore how changes in behavior relate to odor responses, we compared various aspects of odor-related PC population activity (**Figure 3**) to our behavioral score. We first compare changes in overall mean odor response magnitude for all neurons and odors. Regardless of day, we found that mean odor response magnitude closely followed the odor-evoked behavior of the animals with each decreasing across trials and rebounding on the following day (Day1: R^2^= 0.86; p = 0.001, Day2: R^2^ = 0.76; p = 0.005, Day3: R^2^ = 0.90; p = 0.001, Day4: R^2^ = 0.79; p = 0.003, Day5: R^2^= 0.87; p = 0.001) (**Figure 8D**). As with response magnitude, changes in the percent of neurons significantly responding within a day closely followed the odor-evoked behavior of the animals (Day1: R^2^= 0.87; p = 0.001, Day2: R^2^= 0.74; p = 0.006, Day3: R^2^= 0.94; p = 0.001, Day4: R^2^= 0.96; 0.001, Day5: R^2^ = 0.95; p = 0.001) **(Figure 8E**). Interestingly, when including only neurons that were significantly responding to each odor, we found little relationship to the odor-evoked behavior of the animals (Day1 R^2^ = 0.41; p = 0.09, Day2 R^2^= 0.35; p = 0.12, Day3 R^2^= 0.69; p = 0.01, Day4 R^2^= 0.03; p = 0.703, Day5 R^2^= 0.28; p = 0.001) (**Figure 8F**). In total, this provided evidence that investigative behavior may influence overall aPC activity independent of odor response profile. Given prior evidence for decreases in OB output in response to repeated odor experience (Kato et al., 2012; Ross and Fletcher, 2018), the observed decrease in stronger responses to odor presentation may be more associated with experience-dependent plasticity mechanisms rather than behavior.

Finally, given the relationship between overall PC activity and the behavioral score we observed above, we tested whether the investigatory behavior of the animal could explain any of the variability we observed in response consistency across days. Given that the most consistent representation of odor identity was found within the first day of recording (**Figure 4C**), we focused on the consistency of responses on Days 2-5 when compared back to Day 1. Specifically, we compared the correlation of each trial on days 2-5 to each trial on day 1 for the best odor from each animal (**Figure 9A**). There was a significant decrease in correlation across trials (2-way ANOVA, main effect of trial, F(7, 224) = 130.2, p < 0.001) independent of the decrease in cross-day consistency (2-way ANOVA, main effect of trial, F(3, 224) = 119.3, p < 0.001) (**Figure 9B**). Post hoc tests comparing trial 1 to trial 8 revealed this effect was significant for all days (p < 0.001). To compare this to behavior, we plotted each trial’s mean correlation to all Day 1 trials versus the mean behavior score elicited by that odor. Overall, when pooling all days, we found a significant positive relationship between the two (R^2^ = 0.51; p < 0.001), showing that trials that were highly correlated to Day 1 responses also showed higher behavior scores (**Figure 9C**). To further test this independently of our sorting for the best-represented odor, we also trained a classifier to predict odor identity based on the behavioral score of each trial. We split responses across Days 2-5 into two groups, with one group containing the responses from the two trials with the highest behavioral score on each day and the other containing the responses from the two trials with the lowest behavioral score on each day. This allowed for a similar model construction to prior figures, as the same trial numbers (8 for each odor) were used as in prior figures (**Figure 4,5**). Compared to the data from higher behavioral score trials, we found classifier performance to be substantially worse in decoding odor identity when using the lower behavioral score trials (**Figure 9D**). In summation, this provides evidence that behavioral state impacts overall activity in aPC as well as the consistency and reliability of long-term representation of odor identity.

**Figure 9.**
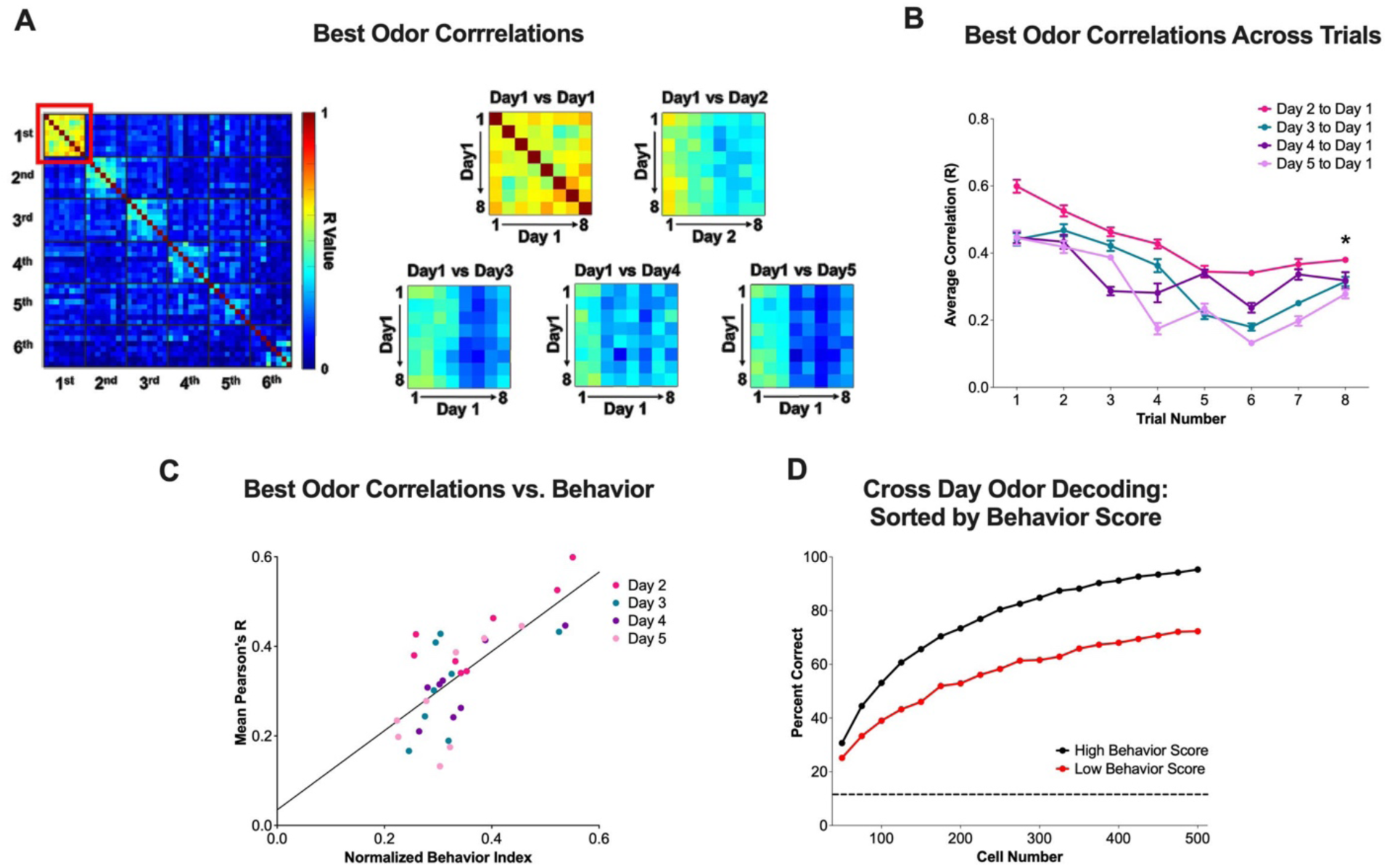
Cross-day odor response consistency is impacted by behavior. **(A)** Highlighting previous comparison of trial-to-trial correlations to Day 1 when sorted by the best odor (Figure 5B); emphasis on the change in response consistency across time within a session for the best odor. **(B)** Quantification of correlations to Day 1 by trial number for the best odor as shown in A; mean and SEM shown. * denotes a significant difference from Trial 1. **(C)** Correlation of normalized behavior index (from Figure 8C) on Days 2-5 to the mean correlation to Day 1 of the best odor on the same days shown in B. Regression line of best fit shown for all days. **D)** Odor decoding from increasingly large random subsets of the neural data using an LDA classifier on data from Days 2-5. For each odor on each day, trials were sorted by their behavior index values on that trial and separated into two new datasets: a “High” score dataset containing the two trials of each odor on each day with the largest behavior index values and a “Low” score dataset containing the two trials of each odor on each day with the lowest behavior index values. LDA models were then trained/tested using leave-one-out validation using only the “High” or “Low” score data (equivalent trial numbers used to train each model to previous figures). For each data point, 100 iterations of randomly sampled neurons from the dataset of size n=neuron number were used to train each model. The same neurons were used across days at each iteration for consistency.

## Discussion

Here, we present a novel dataset characterizing activity from the same population of aPC neurons over multiple days from freely moving mice receiving passive presentations of odor stimuli. Overall, we recorded dynamic behavior during these sessions, with animals demonstrating both behavioral habituation within sessions alongside greater investigation at the beginning of each session. This allowed us to demonstrate that overall activity in aPC closely correlates with investigatory behavior, similar to previous work showing decreases in population activity with changing sniff rate (Bolding et al., 2020). Despite the significant differences in our experiments from previous head-fixed odor presentation paradigms, we also found that odor-trial evoked ensemble response within a day represented odor identity, recapitulating prior results (Bolding and Franks, 2018, 2017; Rennaker et al., 2007; Roland et al., 2017; Stettler and Axel, 2009), but that these ensembles decrease in similarity over time. The neurons comprising the ensembles that best represent odors on a given day are not 1:1 consistent across time, similar to recent evidence suggesting representational drift in aPC (Schoonover et al., 2021). Importantly, our behavioral measures indicate that cross-day odor response consistency correlates with odor investigation. Combined, this indicates two interacting effects that influence aPC odor coding: experience-dependent decreases in the robustness of odor responses alongside a decrease in cross-day ensemble consistency that is modulated by behavioral state.

Prior acute recordings established a role for aPC responses mediating sensory habituation, with repeated presentations of odors driving decreases in the output of individual neurons linked to identity representation (Kadohisa and Wilson, 2006; Wilson, 2000, 1998). Despite the fact that our experiments were designed specifically without repeats of any given odors, we found substantial habituation both in terms of response magnitude and percent of responsive neurons. Our observed changes in response magnitude fit well with prior studies recording longitudinally in the OB that have shown habituation of odor-evoked glomerular and M/T responses with passive odor experience (Kato et al., 2012; Ross and Fletcher, 2018). While changes in cross-day response similarity appear to stem from a reorganization of neuron response profile (**Figure 7**), decreases in OB output to aPC may still be driving at least some component of the observed decreases in within-session response similarity (**Figure 4**). Subsequent studies exploring these two effects may clarify their relationship. Additionally, learning can recover or enhance OB responses to the same odor stimulus (Kass and McGann, 2017; Ross and Fletcher, 2018), possibly explaining the maintained consistency of responses observed in aPC studies where animals are performing tasks (P. Y. Wang et al., 2020).

Two recent studies have also tracked a consistent population of aPC neurons over multiple days of recording odor responses and provided the most direct comparisons to this work, though both used head-fixed odor presentation paradigms. One collected electrophysiological recordings from a population of aPC neurons across longer periods of experience (Schoonover et al., 2021). They found significant representational drift in aPC when animals only received intermittent odor experience, and that drift rate could be reduced by frequent experience. While it is difficult to directly compare our data to this study due to the different recording techniques, stimulus delivery methods, and number of trials within a session, our data does support a degree of inconsistency in aPC responses over time, even with more frequent experience. We still found some overlap between aPC responses from Day 1 and Day 5 (**Figure 5**), indicating that ensemble shifts found in our data were certainly not a 100% reorganization of aPC odor representations. Follow-up freely-moving recordings with less frequent experience may provide a more direct comparison between these studies, as non-continuous experience may result in similar complete shifts in response ensemble observed in Schoonover et al. In a second study, responses were recorded before and after an appetitive learning paradigm (P. Y. Wang et al., 2020) by 2-photon microscopy, finding minimal changes in aPC activity. The authors reported modest enhancements in paired odor response ensemble similarity and classification accuracy after learning relative to controls but no other major changes, focusing on downstream targets of aPC for the remainder of the study. We would attribute the differential effect of odor experience, decreasing classification accuracy and response correlation in our data, to differences in behavioral paradigms. In our study, animals were allowed to freely sample odors, giving or withholding attention, while the head-fixed paradigm reported by Wang et al. would demand focus from the animals as they were performing an olfactory-driven task. Additionally, we are also not reporting any associative learning paradigms in the data presented here, and there remains the possibility that the decreases in response correlation we observed could be attenuated by a similar appetitive learning paradigm.

Our observed link between investigative behavior and decreased response similarity provides an explanation for some of the response ensemble inconsistencies we found across days. Behavioral modulation of cortical activity is a potential confounding factor in a number of studies assessing representational consistency, as animal attention directly impacts the sampling of a stimulus (Sadeh and Clopath, 2022). Imaging studies in the human olfactory system have indicated a role in attentional modulation of PC function (Arabkheradmand et al., 2020; Zelano et al., 2005), and work in rodents has demonstrated an impact of the behavioral state on the olfactory network (Carlson et al., 2018; Schreck et al., 2022). By our behavioral metrics, within-day shifts in odor investigation may impact coding similarity across days (**Figure 9**); that said, our behavioral measures do not account for all of the shifts in ensemble coding. This implies the possibility of complementary mechanisms that could underly response ensemble inconsistency; long-term plasticity mechanisms within aPC may reorganize coding ensembles over time (drift), and behavioral focus may modulate reliability on shorter timescales. While our behavioral readouts give us relative assessments of attention across trials, they do not provide absolute readouts of brain state or arousal. Future experiments may be able to leverage direct readouts of attention, such as sniff rate (Bolding et al., 2020; Ogg et al., 2018) or pupil dilation (Montes-Lourido et al., 2021) to compile more direct measures of animal attention. Anecdotally, we also observed that many odor-driven behaviors, such as grooming, were able to activate aPC as strongly as our panel of odors. Categorizing these behavioral events was not the primary goal of this study but will likely be addressed in future work, particularly given the link between control of grooming and other regions of the olfactory cortex (OC) (Zhang et al., 2021). In total, this work reinforces prior studies indicating a role for aPC in the formation of odor representations, despite the use of a new, freely-moving odor sampling paradigm but demonstrates a role for both experience and behavioral state in shaping the consistency of these representations over time.

While the miniscope technique provided the unprecedented ability to track conserved cellular activity in aPC, implementation of these recordings in freely-moving animals did introduce technical limitations that should be kept in mind when comparing our results to prior studies. First, while there are functionally distinct cell types in aPC possibly contributing differentially to these odor responses (Diodato et al., 2016; Nagappan and Franks, 2021; Suzuki and Bekkers, 2011; L. Wang et al., 2020), the miniscope recording technique integrates from a large imaging z-plane, not allowing us to clearly distinguish between cell types and layers of aPC as with 2-photon imaging. For the sake of this report, we, therefore, chose to focus on population-level responses from the network as a whole. Primary components of our results, such as response correlation and lifetime sparseness, generally match those collected with awake recording techniques (Bolding et al., 2020; Bolding and Franks, 2017; Miura et al., 2012; Schoonover et al., 2021). Recording in freely moving and uncued animals likely provided another source of variation; while it provided a previously unexplored study of naturalistic odor sampling, we do not have the same degree of stimulus control as in head-fixed recordings. Due to these drawbacks, we focused our analyses on population ensemble response properties instead of timing encoded information; there remains the possibility that a subset of more quickly responsive neurons shown in past primacy coding experiments may maintain greater consistency in responses across days (Bolding and Franks, 2017; Wilson et al., 2017).

## Acknowledgments

We thank Brittany Correia for her help with the analysis code.

## Author contributions

I.F.C. and M.L.F. conceptualized the project and designed the experiments. I.F.C. conducted the surgery, lens implantation, and data collection. I.F.C. and M.A.R. processed and analyzed the data. I.F.C. and M.L.F. wrote the manuscript with input from M.A.R.

## Methods

### Animals

All experiments were approved by the UTHSC Institutional Animal Care and Use Committee. Both male and female (n=16 total) C56BL/6J mice were used for all experiments; animals were approximately 12 weeks of age when we began surgical procedures. Mice were housed on a standard 12:12 day/night cycle, with all recordings collected during their light cycle (6 AM-6 PM). All animals were individually housed after GRIN lens implantation. Animals were excluded from the dataset if the integrity of their imaging fields degraded visibly throughout the experiment or if the imaging Z-plane shifted dramatically across days. Males and females within the given age range were equally capable of carrying the weight of the miniscope throughout recording sessions.

### Virus Injection and Lens Implantation

All surgical procedures began at approximately 12 weeks of age. Mice were anesthetized by isoflurane (5% induction/2% maintenance) for all surgical procedures and were given pre-operative analgesics (carprofen 5 mg/kg) subcutaneously. For virus surgeries, animals were secured in a stereotaxic apparatus (David Kopf Instruments), and an incision was made to locate the Bregma and Lamba sutures. Hydrogen peroxide (0.3%) was applied to debride the skull and expose the sutures. An approximately 0.5 mm diameter craniotomy was made using a dental drill (Narashige) +1.7 mm anterior and +2.3 mm lateral from Bregma on the right side of the skull. The injector (Drummond Nanoject III) needle was lowered 3.3 mm from the surface of the brain to target aPC. The pipette was left in place at the target site for 10 minutes prior to injection to allow any displaced tissue to settle, and then 500 nL of AAV1-hSyn-GCaMP6s-WPRE-SV40 (Addgene: 100843-AAV1) was injected at a rate of 5 nL/second. After allowing 10 minutes for virus diffusion, the injector was slowly retracted from the brain, the skin was sutured, and animals were allowed to recover for 2 weeks before later GRIN lens implantation.

For GRIN lens implantation, animals were anesthetized and secured in the stereotaxic apparatus as before. The skull was gently scored with a scalpel, and a screw was implanted in the left parietal bone to strengthen the subsequent head cap. A craniotomy was made in the same location as the virus injection of approximately 0.6 mm in diameter to accommodate the GRIN lens, and a custom vacuum system held the GRIN lens in place as it was lowered into the craniotomy. For every 1 mm of depth lowered, the lens was allowed to rest for 5 minutes to minimize damage to surrounding tissue. Once the lens was lowered to 3.3 mm from the surface of the brain, it was secured in place with cyanoacrylic glue (Bob Smith Industries). The surface of the skull was then covered with a thin layer of the same glue, followed by acrylic dental cement (Ortho-Jet). The lens was covered by a thin layer of Kwik-Sil (WPI) and an additional thin layer of dental acrylic to protect the lens surface while the animals recovered from the surgery. Animals were given Carprofen (5 mg/kg) and Dexamethasone (1mg/kg) injections subcutaneously for one week post-operatively to aid with recovery and clearance of blood from around the lens. For some animals, virus injection and GRIN lens implantation were combined into a single surgery, with lens implantation proceeding immediately after virus injection at the same site. In either case, imaging did not begin until at least 6 weeks after the implant of the lens to allow for animal recovery and healing of the imaging ROI.

### Baseplating

Animals were anesthetized and secured in the stereotaxic apparatus as before, and a dental drill was used to remove the dental cement and Kwik-Sil cover from the top of the GRIN lens. The surface of the lens was gently cleaned with ethanol to remove any debris before affixing the baseplate (miniscopeparts.com). A baseplate with a miniscope attached for viewing the ROI was held above the lens using a custom 3-D printed holder. The baseplate was lowered to the ROI at the plane where neurons and landmarks such as blood vessels were most clearly resolved. The baseplate was initially affixed with cyanoacrylate glue, then further secured with a layer of dental cement to minimize ambient light leakage into the miniscope field of view during imaging.

### Histology

After experiments were completed, animals were anesthetized with a ketamine/xylazine mixture (100/10 mg/kg) for trans-cardial perfusion. Animals were perfused with 0.9% w/v saline to remove blood, followed by formalin (4% formaldehyde) to fix the brain. Brains were post-fixed in the skull for one week after perfusion to preserve the shape of the lens track. Brains were then washed in PBS to remove formalin, cryoprotected in 30% w/v sucrose, and cut into 40 µm coronal sections using a freezing microtome. Sections were cover-slipped with Vectashield Plus antifade mounting medium (Vector Laboratories) and imaged using a Nikon Eclipse 90i fluorescent microscope to verify lens placement in aPC.

### Behavioral and Imaging Equipment

All imaging was performed in a two-chambered box where each individual chamber was 9 cm in height and 12.5 cm in diameter. The top of the chamber was loosely covered by a sheet of hard plastic containing a 1 cm diameter hole for the miniscope wire. Animals were restricted to one side of the chamber by a perforated wall between the two chambers. The odor delivery port was affixed to the top of the wall of the chamber occupied by the mouse, while a vacuum line for clearing the odorants was attached to the far wall of the second chamber. This was to avoid any aversion of the animal to the pull from the vacuum line during recording sessions. All recordings were collected under low light conditions in an isolation chamber (Coulbourn Instruments). Acclimation to the miniscope was performed in a clean housing cage prior to any imaging.

A UCLA v4 miniscope (https://github.com/Aharoni-Lab/Miniscope-v4) was used for all data collection. The miniscope was connected to a DAQ for recording (UCLA Miniscope) via a passive commutator (ONE Core University of Colorado) to allow animals to move freely during recording sessions. Framerate, Gain, and LED power were adjusted for each mouse to maximize dynamic response range but were kept consistent across recording days. An IR camera (SenTech) recorded animal behavior during odor presentations from a side view angle at approximately 60 Hz. All miniscope and behavioral recordings were collected through software written for the UCLA miniscope DAQ (https://github.com/Aharoni-Lab/Miniscope-DAQ-QT-Software/wiki) to maintain consistent timestamps.

### Odor Panel and Delivery

For all experiments, the stimulus panel was composed of the following 6 odorants: Benzaldehyde, Butyraldehyde, Ethyl Butyrate, 2-Heptanone, Linalool, and 2,5-Dimethylpyrazine (> 98% purity stocks, all from Sigma-Aldrich). Odors were diluted to 1% weight/volume in light mineral oil (Sigma-Aldrich) in scintillation vials with 18-gauge needles inserted for airflow. A custom olfactometer was built that supplied air from an aquarium pump at a constant rate of 1 liter per minute (LPM). Vials containing odors were connected to individual odor lines that had pinch valves upstream of the vial to close airflow from the aquarium pump when not in use. When not presenting an odor, the air was routed through a clean air line in the odor port, and the vacuum line was on at a matching one LPM flow rate. For odor presentations, a pinch valve would close the clean air line, and the pinch valve along the dedicated line for the specified odor was opened for 5 seconds, allowing odorized air to flow from the scintillation vial into the chamber. The vacuum line was also closed for the duration of the presentation via a separate pinch valve wired to the same switch. Presentation order was pseudorandomized such that no trials of the same odor were repeated, but that trial order was different for each animal on each day. Regardless of presentation day, all trials had a variable inter-trial interval of 40-80 seconds.

### Experiment Schedule

Animals were first acclimated to the weight of the miniscope during 3 days of 10-minute sessions in a clean cage. These sessions were also used to fine-tune imaging settings to best capture the most neurons in the imaging ROI. On the first day following acclimation, animals began the odor-presentation paradigm. For each day of odor presentations, animals were gently restrained, and the miniscope was affixed to their baseplate. The recording ROI was verified to match the expected ROI from acclimation sessions, and then animals were placed in the two-chambered box for odor presentations. Each day, the animals were first acclimated to the box for 10 minutes, then the 10 trials of each odor were presented, followed by another 10-minute period with no odor presentations. At the end of each session, the miniscope was removed, and animals were returned to their homecage. Miniscope and behavioral videos were recorded continuously throughout the session, with timestamps recorded through the DAQ software for each odor presentation.

### Calcium Trace Extraction

For each session, calcium recordings were concatenated into a single TIFF stack. Then, this stack was spatially downsampled by 2x in both the X and Y dimensions and temporally downsampled to 5 Hz. Stacks were cropped to remove the lens edge to minimize processing time when possible. ROIs and traces were then extracted using the Python implementation of CaImAn (Giovannucci et al., 2019). In brief, CaImAn combines NoRMCorre, a motion correction algorithm (Pnevmatikakis and Giovannucci, 2017), and a constrained nonnegative matrix factorization-based ROI detection package (Zhou et al., 2018) into a single pipeline to identify spatial footprints of neurons and calculate temporal components of those neurons. Inputs regarding the size and shape of expected neurons were kept constant for all recording sessions. The minimum signal-to-noise (min_pnr: > 4x background) and minimal spatial correlation (min_corr: 0.9-0.98) values were adjusted for each session but were within the range given above to minimize possible detection of the same neuron as multiple ROIs. A “raw” trace output, where a denoised signal was added back to the deconvolved trace to better represent noise within the recording, was used to assess temporal components of neuronal activity in downstream analyses.

### Neuron Alignment

Neuron footprints extracted by CaImAn were used for alignment with the Cell-Reg package (Sheintuch et al., 2017). Initial alignment was performed, allowing for both translations and rotations up to 15 degrees. Any animals with apparent discrepancies between recording sessions were excluded from analysis, as this would represent significant shifts in the ROI recorded on those days. Final alignment was performed using probabilistic modeling recommended by the Cell-Reg package. Alignment indices were used to focus downstream analysis only on neurons that were identified in all days of miniscope recordings.

### Response Extraction

Custom MATLAB scripts were written to calculate calcium responses from odor presentations. Given that the output of CaImAn represents a background-scaled fluorescence value, a single value (ΔF) was calculated relative to the odor presentation time points to represent the value of a response at that time point. In each case, a given ΔF value was calculated by subtracting the mean value in a 2s window before a time point from the maximal value in a 5-second window after the same time point. For shifted response window analysis (**Figure 2**), all possible ΔF values were initially calculated using all possible time points within a set window from the start of odor onset. For each odor, all possible shifted time points within that window on a given day were compared using the Pearson correlation coefficient. The timepoint on each trial with the maximal R-value (“shifted timepoint”) was used as that trial’s response timepoint in downstream analysis to focus on the point at which PC best represented a given odor during a trial. We used a 7s window to cover the entire odor presentation as well as a short time after odor offset; a longer 10s window was used to demonstrate the distribution of time-to-peak trial-to-trial correlation (**Figure 2**). For comparisons to time points before the odor presentation (**Figure 2**), we used the same shifting window approach on data starting 15 seconds before odor onset to avoid any potential overlap with odor trials. Pre-window pseudo-trials were randomly selected independent of the odor that was preceding or following the pre-trial timepoint to minimize the effect of any potential lingering odor responsivity in the window.

To account for the loss of some trials due to recording or presentation errors, we curated the response dataset to the first 8 kept presentations of each odor on each day. This led to the exclusion of two animals as they did not meet this criterion for trial number on at least one day of recording, trimming the final neural dataset to n=687 neurons tracked across all sessions from n=14 animals. Significantly odor-responsive trials for a neuron were defined as having a post-timepoint response exceeding the pre-timepoint value by at least 2.5 standard deviations, as well as having a total trace z-score value at that maximal point that was > 1. For bulk response characteristics (mean-evoked ΔF, percent responders), trials were averaged across all odors by their relative presentation number (i.e., 1-8 after adjustment) to minimize the impact of the variability in odor response preference from a given animal **(Figure S2)**. For calculating the “best” odor on an animal-to-animal basis (**Figure 4**), odors were re-sorted by the largest mean within the odor trial R-value on the first day of recording.

### Behavioral Pose Estimation

Pose estimation was performed by training a model with DeepLabCut (Mathis et al., 2018; Nath et al., 2019) to extract key-point values from the animal for the generation of kinematic measures of behavior. Behavior videos were first concatenated into a single .mp4 file for each recording session. Sample videos containing animals from different recording days were compiled to serve as training videos for the model and were annotated to identify the following ROIs: base of the tail, center of the body, left ear, right ear, nose, and the body of the miniscope. Annotated frames were used to train a model to approximately 250,000 iterations, at which point training did not lead to an appreciable improvement in model accuracy. Poses from all remaining videos from n=15 animals were then extracted using the trained model. Neural data was recorded from one animal without the side-view camera; therefore, it was not included in the behavioral dataset.

Two primary behavioral measures were calculated using the poses estimated by the DeepLabCut model in a 5s window after the shifted time point indicated during response extraction (same window as neural data for comparison). First, a general motion index was calculated by averaging the Euclidean distance traveled by each tracked ROI between frames. While not specific to any one body part, this index represents the relative activity of the animal. Second, head raising, a stereotypic behavior in mice associated with investigation, was calculated by subtracting the Y coordinate of an ear from the Y coordinate of the nose. When possible, the left ear was used to avoid displacement differences between the two ears. For both measures, a strict confidence cutoff of 0.95 was used in deciding which tracked components to include for each measure. Since behavior was recorded from the side, this cutoff was chosen to avoid including ROIs at time points when tracked points moved in or out of view of the recording camera. The raw values for each metric were calculated by taking the mean value (motion) or mean value +/- six frames around the maximal value (nose) during the window used for calculating neural responses on a given trial. Once extracted, both measures were normalized by dividing by the maximal value of that measure for a given animal on all recording days. Then, the two normalized values were averaged to generate a single behavioral index for relative investigative behavior that ranged between 0 and 1 for all animals.

### Linear Classifier

Classifier analysis was performed using the Statistics and Machine Learning Toolbox in MATLAB. Vectors for each trial of each odor were created by combining all ΔF responses from all neurons tracked across all days of odor presentations. Random sub-populations of neurons of increasing size from the dataset were then selected for training the model, with 100 iterations of random neuron selection performed at each population size step. Leave-one-out validation was then used for training and assessing linear discriminant (LDA) models, i.e., for a given trial, a model was trained using all trials on a given day except for the trial to be tested, and then the excluded trial was tested against the model. This method was then repeated for every trial, and total accuracy was determined by the percentage of correctly classified trials.

### Lifetime Sparseness and Tuning

We chose lifetime sparseness as a measure of tuning as it allows for the assessment of relative responses to different stimuli without having to introduce arbitrary response cutoffs and has been used in prior olfactory coding studies (Bolding and Franks, 2017; Poo and Isaacson, 2009; Roland et al., 2017; Rolls and Tovee, 1995; Schoonover et al., 2021). Lifetime sparseness was calculated using the following formula:

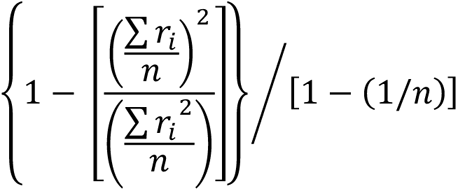

Where *ri* is the summarized response to 1 through *n* total stimuli. For our data, we chose to use a median value for *r* for each stimulus to minimize the effect of high outlier response values that could skew a mean measure for *r*. The lifetime sparseness value for each neuron was then used to assess relative tuning, i.e., neurons with higher lifetime sparseness exhibited more narrow tuning to specific stimuli than those with lower lifetime sparseness values. “Best” odor response tuning for individual neurons on a given day was calculated by sorting neurons by the odor for which they had the highest median response on that day. The top 20% of sparsely responsive neurons, roughly representative of the percent of neurons responding to odor presentations (**Figure 3C, D)**, was used to avoid including neurons that had relatively non-differential responses to odors in the panel on a given day. The average response across trials of every odor was then calculated after sorting the neurons by their best response.

### Statistical Tests

Statistical testing and figure generation were performed in MATLAB or GraphPad Prism. For calculating response correlation, Pearson’s correlation coefficient was calculated using all ΔF values from selected cells for each odor trial. For across-group comparisons, unless otherwise stated, ANOVA with Greenhouse-Geisser corrections and planned post-hoc tests (Fishers LSD) were used to assess significant changes in response magnitude, response correlation, or changes in behavioral readouts.

---

**Supplemental Figure 1.**
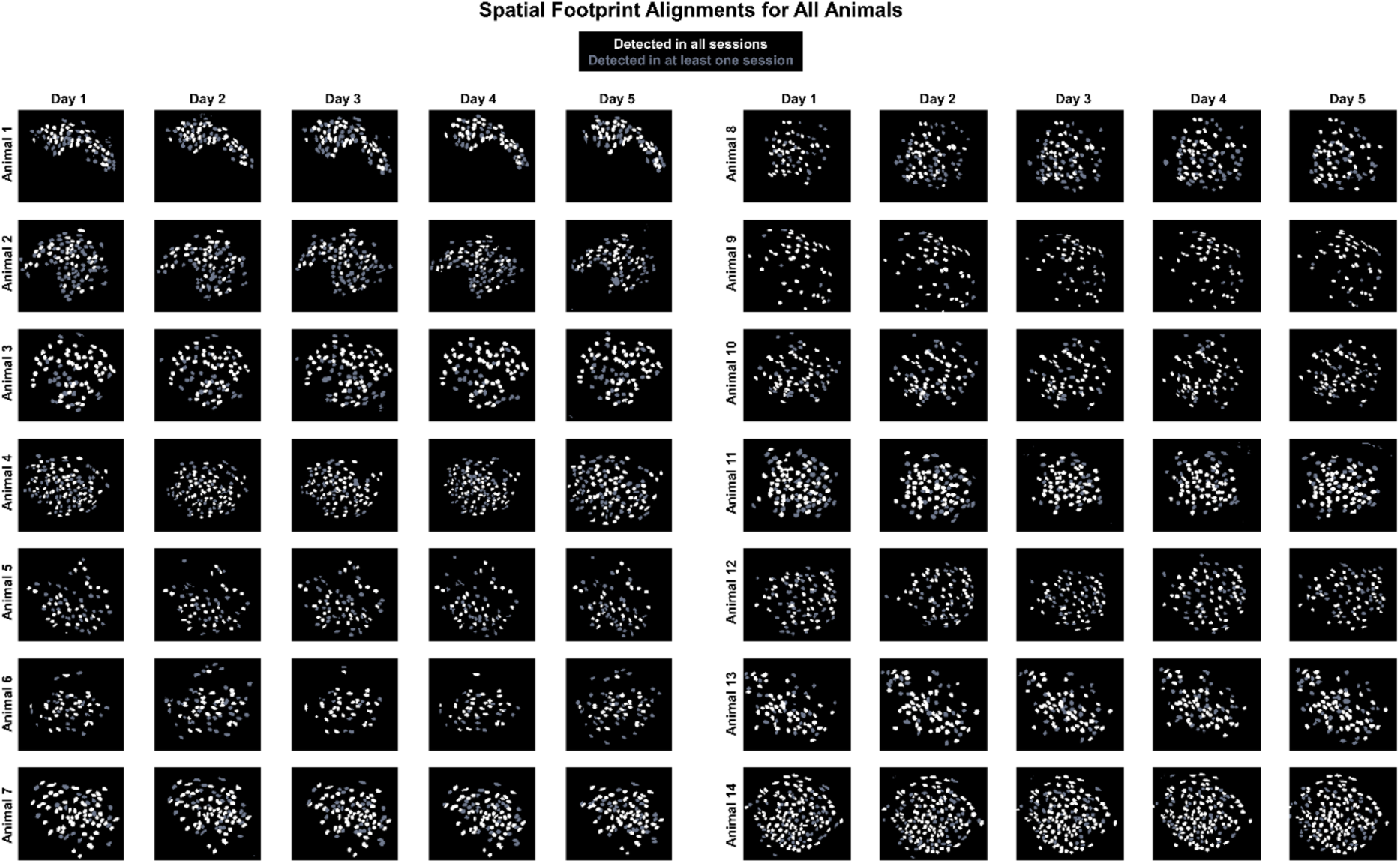
Spatial footprints of neurons aligned across sessions for all animals. Aligned spatial footprints from every recording session for each animal. Grey neurons were detected in at least one session but not all 5 recording sessions, whereas white neurons were identified in all 5 sessions and were therefore used in downstream data analysis.

**Supplemental Figure S2.**
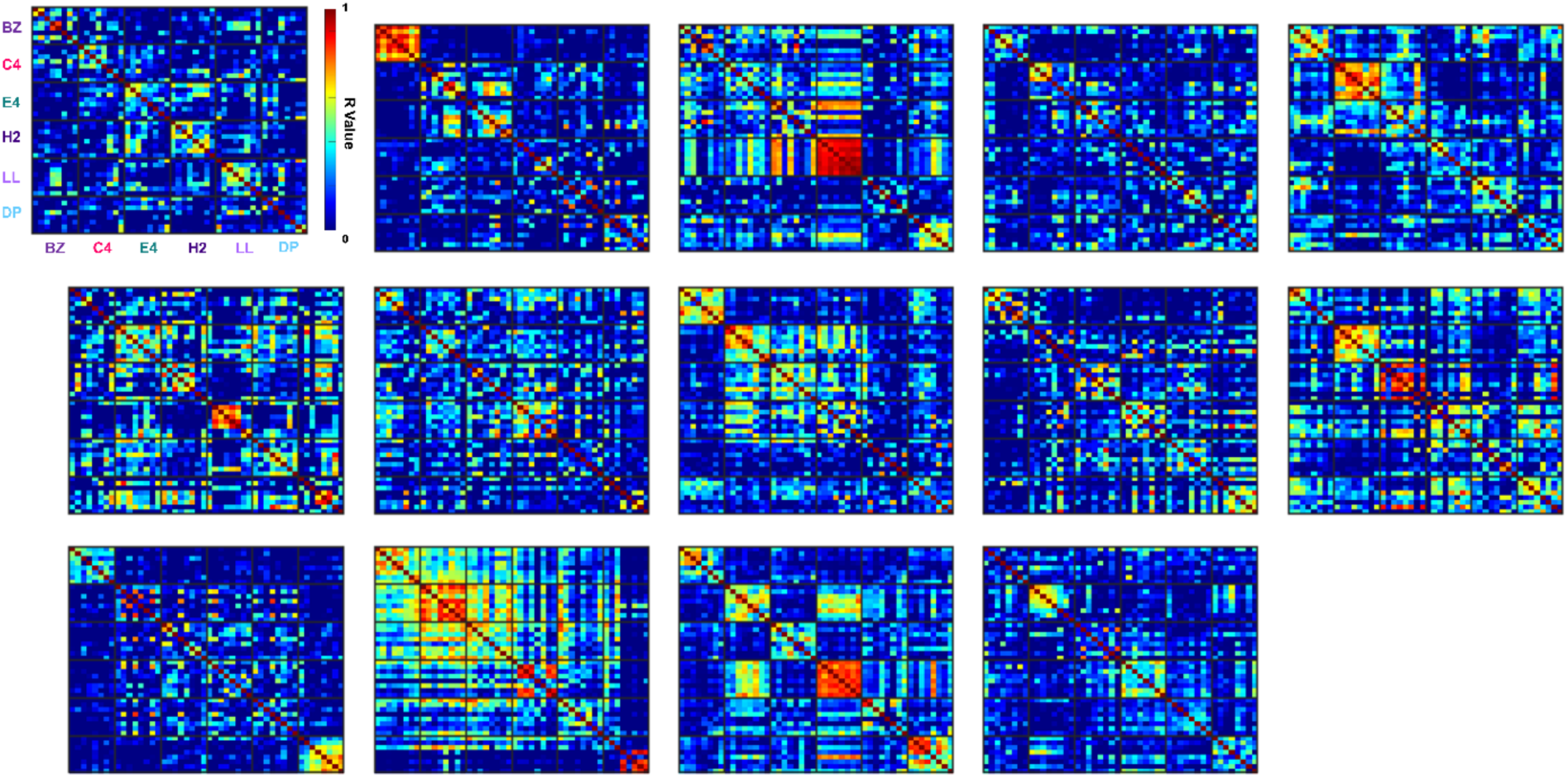
Trial correlation matrices for individual animals. Trial-to-trial correlation (Pearson’s R) matrices were generated between all trials for all neurons on Day 1 of odor presentations; trials were organized by odor identity. Each sub-panel represents all values from an individual animal.

